# Ecosystem disservices are underrepresented in literature on annual crop agroecosystems

**DOI:** 10.64898/2026.06.09.730574

**Authors:** Gábor Pozsgai, Kris A.G. Wyckhuys, Ibtissem Ben Fekih, Liette Vasseur, Geoff Gurr, Mark Goettel, Heather VanVolkenburg, Sunita Pandey, Jamile Queiroz-Sousa, Pingyang Zhu, Mohammed Abul Monjur Khan, Titus Imboma, Margaret M. Hughes, Jian Liu, Syed Rizvi, Olivia Reynolds, Hafiz Sohaib Ahmed Saqib, Stephen Wratten, Jie Zhang, Wenwu Zhou, Francisco Javier Garcia, Lucas Alexander Shuttleworth, Li-Lin Chen, Gábor L. Lövei, Minsheng You

**Author notes:** These authors contributed equally to this work (Gábor Pozsgai, Kris A.G. Wyckhuys, Ibtissem Ben Fekih). Deceased; contributed to this work prior to passing (Stephen Wratten).

## Abstract

Maximising ecosystem service (ES) benefits while minimising ecosystem disservices (EDS) is essential for ecological intensification in annual crop systems. Yet ecosystem disservices are often underreported in the scientific literature, potentially biasing how agroecosystem functioning is understood. To test this, we built a predicted network of plausible links between ecosystem service providers (taxa or functional groups that can deliver ecosystem services or disservices), and the ES or EDS they may provide. We then compared this with a realised network based on systematic Scopus searches of annual crop literature. The predicted network contained 47 nodes and 103 links, whereas the realised network contained 33 nodes and 58 links, representing declines of 29.8% in nodes and 43.7% in links. Overall connectivity declined, especially for highly connected nodes, and four of the nine predicted disservice nodes were absent from the literature. ES links were more likely to be documented than EDS links, and EDS links were three times more likely to be absent. Across all links, ES were reported in 6.6 times more papers than EDS. Projected networks, which map indirect connections by linking ES directly to EDS if they share common providers, showed that these bundled interactions were strongly reduced, obscuring multifunctionality and trade-offs. This systematic underrepresentation of EDS, reflecting a cognitive bias, can inflate perceived benefits, distort the evaluation of key taxa and interactions, and create unrealistic expectations about intervention outcomes in biological crop protection. Addressing EDS alongside ES is therefore essential for more balanced assessments of crop-system management and better-informed decisions.

## Introduction

Agriculture traditionally relies on ecosystem functions and services that provide benefits for humans. Yet, industrialised forms of agriculture negatively impact provisioning, regulating, and cultural ‘ecosystem services’ (ES) (Millennium Ecosystem Assessment (MEA), 2005), and also undermine supporting ES, such as nutrient cycling or primary production (MEA 2005; Díaz et al., 2018). While several of these ES underpin the functional integrity of agroecosystems, they are degraded through monoculture and biodiversity loss, chemical intensification, land-use change, and the decline of the biogeochemical cycles (Schaller et al., 2018). Meanwhile, the latter processes also negatively affect the myriad ES provided directly by agroecosystems.

In response to these pressures, ecological intensification has emerged as a framework that seeks to sustain agricultural production by managing the ecological processes that underpin crop performance, rather than relying predominantly on external inputs (Bommarco et al., 2013). Central to this approach is the deliberate harnessing of ecosystem services such as pollination, biological pest control, nutrient cycling, soil formation, and water regulation through the conservation or management of biodiversity and habitat complexity. Within this context, sustainable forms of agriculture improve ES alongside on- and off-farm biodiversity (Ramírez-Suárez et al., 2024), which is critically important for ES that are anchored in this biodiversity, such as crop pollination or biological pest control (González-Chang et al., 2020; Wyckhuys et al., 2025). However, whilst ES are provided by ecosystem service providers (ESPs), a broad range of organisms, such as crop pests, plant pathogens, or plant competitors (i.e., weeds), also cause ecosystem disservices (EDS), which hamper productivity and overall functionality (Zhang et al., 2007). Hence, to intensify agri-food production in an ecologically sound manner (Bommarco et al., 2013), one needs to optimize ES delivery while reducing EDS (Dunn, 2010; Power, 2010; Shackleton et al., 2016; Schaubroeck, 2017; Campagne et al., 2018). Since this strategy relies upon accurate knowledge of which organisms and under what circumstances they provide beneficial functions, as well as the possible harms and trade-offs they involve (Blanco et al., 2019; Schaubroeck, 2017), it is especially vulnerable to distorted evidence when ecosystem services are more readily documented than ecosystem disservices.

Yet, while ES are widely appreciated and extensively studied, EDS are regularly overlooked, and a 15-fold difference in the number of papers covering ES versus EDS has been reported (Blanco et al., 2019). Even when accounting for certain methodological shortfalls (Lyytimäki, 2014; Lyytimäki and Sipilä, 2009), this disparity remains (Blanco et al., 2019). Beyond simple underrepresentation, uneven documentation of services and disservices linked to organisms can also distort how agroecosystems are interpreted and managed. When certain taxa or interactions are repeatedly framed in beneficial terms, the literature may create an overly optimistic picture of ecological intensification, even where context-dependent costs or trade-offs exist. This is especially relevant in annual crop systems, where farmers make decisions under increased uncertainty and where insufficient attention to disservices can undermine confidence in biodiversity-based management (Billaud et al., 2025). Since some organisms are traditionally considered to be either beneficial or harmful, a strong bias in attributing the value of their contributions to ES versus EDS most likely exists (Kellert, 1993; Kelemen et al., 2013; Cheng et al., 2023). These distortions are likely reinforced by two interacting filters: taxonomic visibility and cognitive framing. Conspicuous or culturally valued organisms are more likely to be studied and interpreted as beneficial, whereas cryptic, belowground, or negatively perceived taxa may be ignored or considered mainly through a disservice lens (Fisher et al., 2011; Martínez-Sastre et al., 2020; Troudet et al., 2017). As a result, the current scientific record may not simply reflect ecological reality, but also the selective attention of researchers, institutions, and funding systems, increasing the chances that during the planning of sustainable interventions, e.g., in crop production systems (De Heij and Willenborg, 2020), the importance of EDS is underestimated. A more balanced understanding of the relative benefits and trade-offs of ES and EDS, bundled through common ESPs, could therefore help to assess the true usefulness of specific organisms in agroecosystems (Gillespie and Wratten, 2017; Tatiana et al., 2022). Because individual taxa often act as multifunctional providers, i.e. simultaneously delivering both services and disservices, evaluating these bundled effects is critical to accurately determining when the same organisms generate trade-offs across functions (e.g. Tschumi et al., 2018).

Here, we argue that the scientific literature does not merely document agroecosystem functioning but selectively renders some interactions visible while leaving others effectively invisible. If ecosystem services (ES) are more readily studied, valued, and reported than ecosystem disservices (EDS), then the published record may present a systematically distorted picture of how taxa contribute to agricultural multifunctionality. To examine this, we first constructed a conceptually predicted network of plausible associations between ecosystem service providers (ESPs; taxonomic groups) and ES or EDS in annual crop agroecosystems, based on expert knowledge. We then systematically evaluated each of these predefined associations through targeted literature searches (Scopus queries combining ESP and ES/EDS terms) to generate a realised network: associations unsupported by any publications were removed, while supported links were weighted by the number of retrieved studies. This approach allowed us to assess the extent to which ES- and EDS-related associations are represented in the literature, identify knowledge gaps, and determine which ecological relationships remain systematically overlooked. To further explore structural patterns, we analysed indirect connections by collapsing the ESP–ES/EDS network into one-mode projected networks, in which ES or EDS nodes (or ESP taxa) are connected if they share association partners, thereby quantifying the degree of linkedness between ES and EDS and among major ESP taxa. We hypothesised that (1) the realised network deviates from the predicted network due to selective research biases, (2) taxonomic biases are present, with certain groups, particularly small, cryptic, or soil-dwelling taxa, being systematically underrepresented, and (3) ES–EDS co-occurrences mediated by the same providers are underrepresented relative to random expectations.

## Materials and methods

Our analytical approach comprised four main steps: (1) construction of a hypothetical network linking ESPs with ES and EDS based on expert knowledge; (2) an initial Scopus search in 2017, followed by manual screening to generate a human-curated dataset for training and benchmarking machine-learning and AI-based filtering methods; (3) a second, updated Scopus search in 2025, which was screened automatically using the best-performing validated model; and (4) statistical network analyses comparing the predicted network with the final realised network derived from the 2025 literature dataset (Fig. 1).

**Figure 1:**
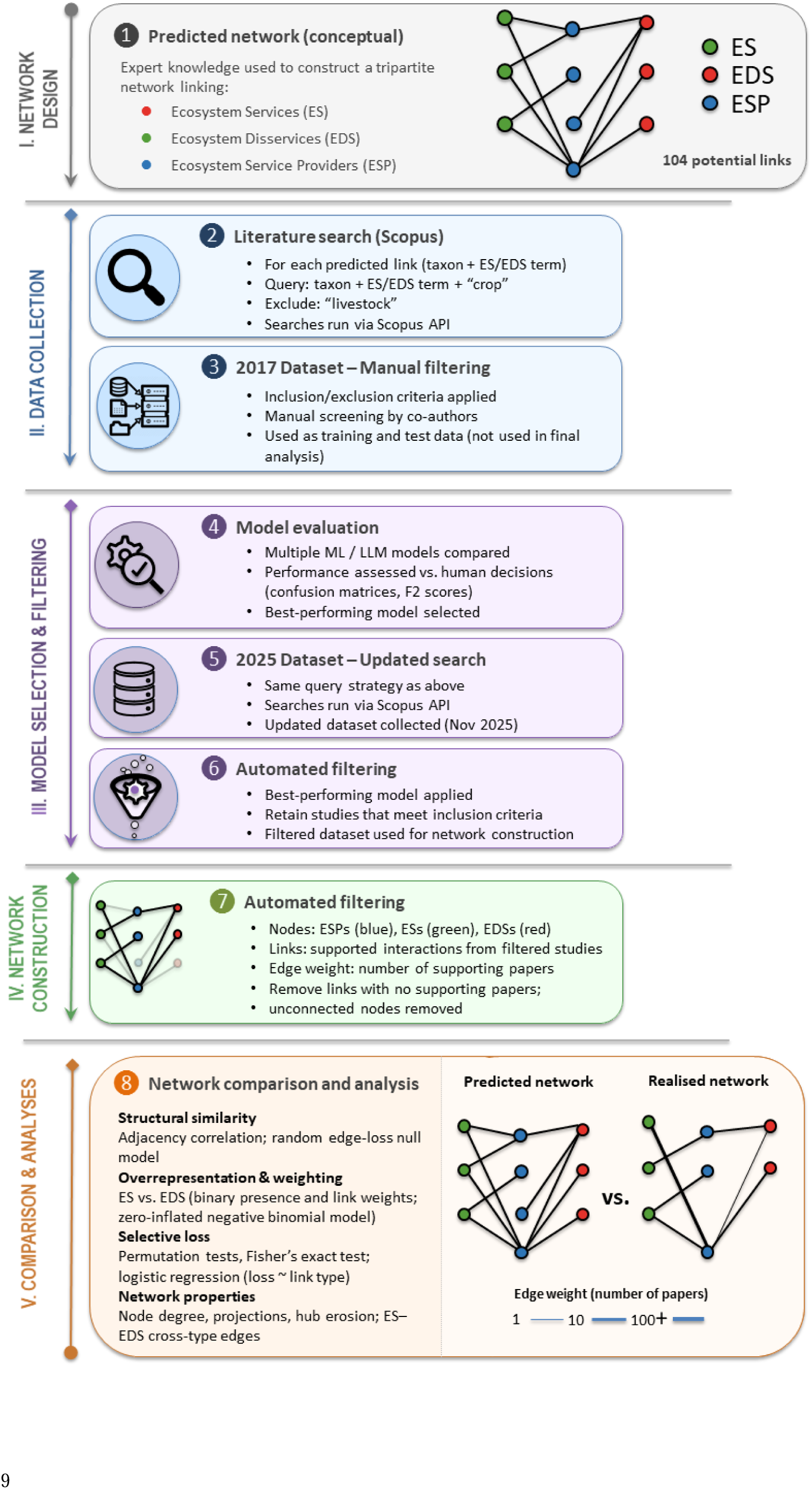
Study workflow. A predicted network linking ecosystem service providers (ESPs) to ecosystem services (ES) and disservices (EDS) was constructed from expert knowledge. Each predicted link was used to query the Scopus database in 2017 and 2025. The 2017 dataset was manually filtered and used to train and evaluate machine learning models, of which the best-performing model was applied to an updated 2025 dataset to automatically identify relevant studies. The retained studies were used to construct a realised, weighted knowledge network of supported associations. Predicted and realised networks were then compared using network statistics.

### Hypothetical network

First, the authors, drawing on their expert knowledge, constructed a hypothetical network indicating probable connections of major plant, animal, and microorganism taxa (ESP) with ecosystem services (ES) and ecosystem disservices (EDS). In doing so, we focused on valid taxonomic units instead of functional groups e.g. pollination whenever possible but did not consider the level of taxonomic organisation (i.e. some taxa may represent families, other classes, or even phyla). This predicted network was used later as a basis for the literature searches (Fig. 2).

**Figure 2.**
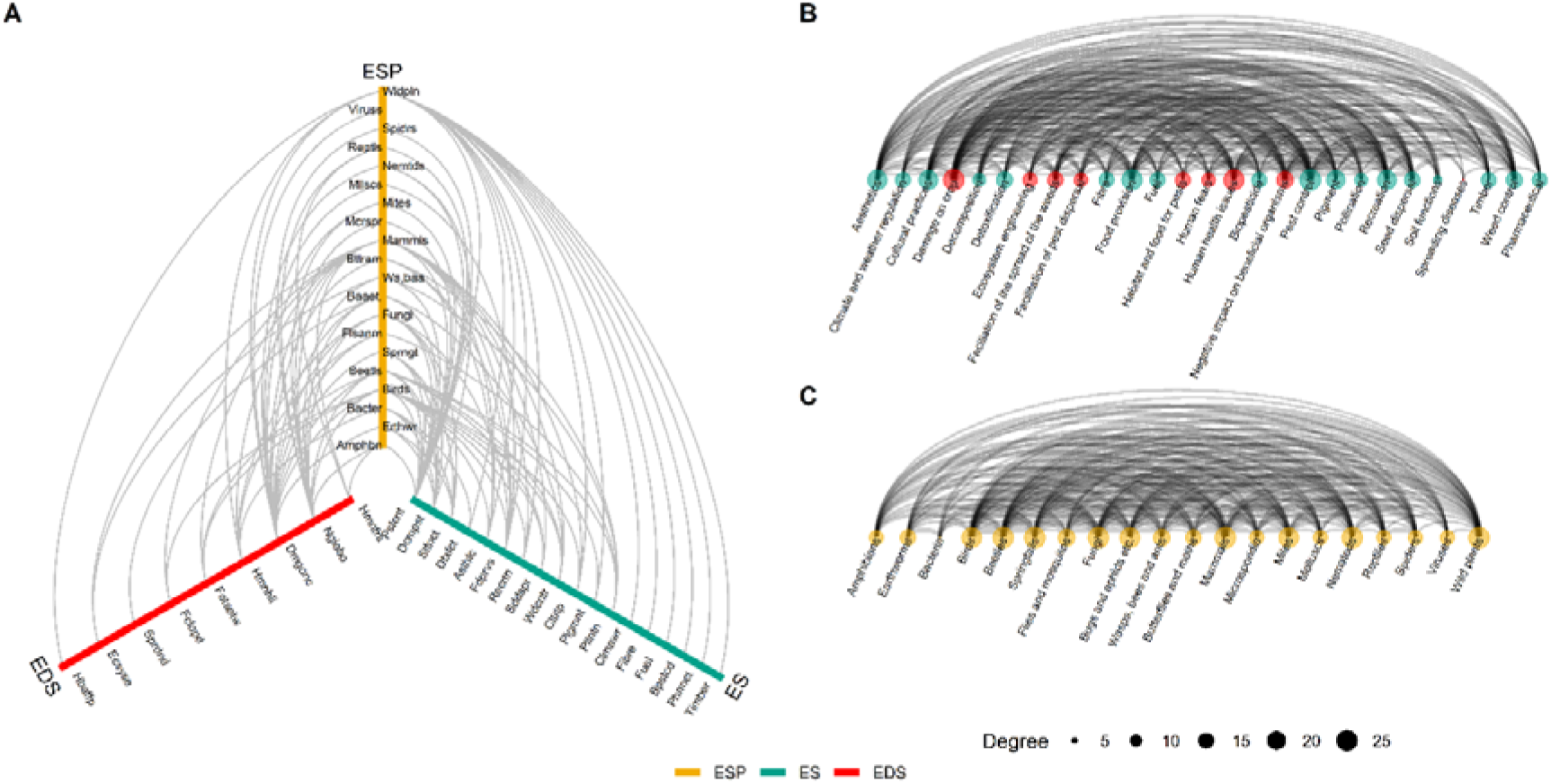
**(A)** Tripartite relationships between ecosystem services (ES), ecosystem disservices (EDS), and ecosystem service providers (ESP) in the predicted network. **(B)** The first projection of the predicted network connecting ES and EDS to each other, based on common links to ESP. Green dots represent ES, red dots EDS. The size of the dots is proportional to the degree of an ES/EDS. **(C)** The second projection of the predicted network connecting ESPs to each other, based on common links to ES/EDS. Size of the dots is proportional to the degree of a particular ESP. Abbreviations are listed in Supplementary Tables 2-4.

### Building realised networks

The respective taxon and ensuing ES or EDS of each of the potential connections between an ESP and an ES or ESD found in the predicted network were used as keywords in a Scopus search. Since our focus was on annual crop systems, we further narrowed the query by adding the word ‘crop’ and excluding the word ‘livestock’. To maximize potential hits, wildcards were periodically included in the Scopus search strings, especially in cases where English common names of plant or animal taxa originated from the scientific names, e.g. mammal* would find mammal, mammals, mammalian. As ‘ecosystem services’ only features in post-1990s literature, we refrained from using this term and instead referred to services or functions such as pollination, nutrient cycling, or biological pest control. This approach was also more feasible because of the underrepresentation of EDS in published literature (Campagne et al., 2018; Shapiro and Báldi, 2014), which would have skewed our foundational dataset. Altogether 104 potential links between ESPs and ES or EDS were traced using customized search strings (Supplementary Table 1).

### Preliminary realised network

First, we ran systematic literature searches from the Charles Sturt University campus in Orange, New South Wales, Australia, on 17^th^ December 2017. Scopus (https://www.scopus.com/) searches were automated using the application programming interface (API) provided by Scopus, to collect published papers where the collective content of title, keywords, and abstract contained the keywords. The API was run under an R environment (R Core Team, 2012).

Papers based on laboratory trials, dealing with non-cash crops, GMO testing, or pesticide efficacy assays, were manually discarded. Reviews, book chapters, or publications other than original research articles or those written in languages other than English were also excluded. Since their inclusion in Scopus’s database was not consistent (https://www.elsevier.com/solutions/scopus/how-scopus-works/content, accessed on 10^th^ November 2025), papers published before 1970 were also not used. The decision to include a paper in the final set was based on the title, keywords, and abstract. This selection process was conducted manually via an online platform that enabled the co-authors to include or exclude papers.

### Final realised network

This foundational dataset was updated through a follow-up search on 11^th^ November, 2025, run through the server of the University of the Azores. Only the newer dataset was used in the final analysis, whereas the 2017 dataset was used to assess the efficiency of machine learning approaches in filtering out irrelevant papers (see below).

While both the 2017 and 2025 datasets were obtained using the same Scopus search procedure, the 2025 dataset was automatically filtered using a machine-learning approach rather than manual screening, based on the same inclusion and exclusion criteria. Before this final filtering, we chose a method that most aligns with human decisions and used the manually filtered 2017 dataset to validate or ground-truth the accuracy of machine learning methods. Machine learning methods ranged from the simple ones, such as penalised logistic regression, random forest, gradient boosted trees, linear support vector machine, and naive Bayes classifier, to advanced large language models (LLMs), such as Open AI’s GPT-5.2 (https://chatgpt.com/), and Perplexity’s Sonar AI (https://www.perplexity.ai), Claude Opus 4.5 (https://claude.ai), and Gemini 3 Pro (https://gemini.google.com). For automating the text classification, we also tested a Qwen2.5□14B LLM, run locally via Ollama (version 3.2, https://ollama.com/) on a workstation with an NVIDIA RTX 5070 GPU and 16 GB VRAM. The model was used in instruction□following chat mode with a maximum context length of 4096 tokens, temperature 0.15, and a maximum of 10 generated tokens per query.

For the non-LLMs, the first dataset containing a unique identifier, the three relevant columns (i.e. title, keywords, and abstract), as well as the human decisions, was randomly split into 80% training and 20% test datasets, and the resulting models were compared using confusion matrices and F2 scores. For LLMs, only the three relevant columns (along with a unique identifier) were provided, the model was prompted to decide to include or exclude papers along the abovementioned criteria, and results were compared to human decisions using confusion matrices, accuracy and precision calculations, as well as F2 scores (Supplementary Figures. 1-10, https://github.com/pozsgaig/cognitive_bias). As the LLM Qwen2.5□14B provided 89% agreement with human decision, it was used to filter the entire 2025 dataset. Only the publications from this filtering were used in the subsequent analysis.

### Statistical analysis

Predicted and realised joint-mention networks between ESP and ES or EDS keywords were represented as graphs. The predicted network captured all theoretically or mechanistically plausible interactions, whereas the realised network represented interactions supported by empirical literature. If the second-phase search did not yield publications for a specific predicted connection, that link and any nodes left unconnected were eliminated. To reflect evidence strength in the realised network, edges between each ESP and ES or EDS were weighted by the number of papers supporting that association. Since the realised network is always a subgraph of the predicted network, all hypothesis tests were evaluated against null models that explicitly accounted for this nested structure.

To test for global structural divergence, both networks were converted to undirected forms and represented as binary adjacency matrices with identical node ordering. Structural similarity was quantified as the Pearson Correlation between the upper triangles of the predicted and realised adjacency matrices. Significance was evaluated using a random edge-loss null model, in which the same number of edges observed in the realised network were randomly removed from the predicted network through 10,000 permutations. The observed adjacency similarity was compared to the resulting null distribution to test whether the realised network was less like the predicted network than expected under random thinning.

Any overrepresented or overweighted ES were quantified using permutation tests in which ES/EDS labels were randomly reassigned among predicted links while preserving the total number of realised interactions. Binary overrepresentation was quantified as the difference in realisation probability between ES and EDS links. Meanwhile, differences in link weights (i.e. the number of publications) were analysed using a zero-inflated negative binomial model, separating (i) the probability that a predicted link was absent from the literature and, (ii) the number of studies supporting realised links. Link type (ES vs EDS) was included as the sole predictor.

To test whether EDS links were more likely to be lost from the predicted to realised networks, we used a permutational approach. All predicted links were classified as ES or EDS and coded as realised or lost based on their presence in the realised network. Selective loss was tested using a permutation test at the link level, in which ES/EDS labels were randomly reassigned among predicted links while preserving the total number of realised links through 10,000 permutations. Next, effect sizes were calculated as the difference in loss probability between EDS and ES links. Using the same label□permutation framework, we also compared ES versus EDS realisation probabilities, defined as the probability that a predicted link appears in the realised network. For both realisation and loss, we computed exact p□values from 2×2 contingency tables (ES vs EDS × realised vs lost) using Fisher’s exact test. Finally, we fitted a logistic regression model with link loss (lost vs realised) as the binary response and link type (EDS vs ES) as the predictor to obtain odds ratios and 95% confidence intervals for the relative loss risk of EDS versus ES links.

The networks were also compared based on the main indices used to characterize networks. Node-level connectivity was quantified as node degree (i.e. the number of links a node has to other nodes) in predicted and realised networks, and the most connected nodes were selected for the ESPs, ES, and EDS in each network. For both bipartite networks (i.e. ESP on one side and ES or EDS on the other side), unipartite side networks (i.e. either ESP or ES/EDS in the network, no clearly separable sides) were projected for both sides. This process connected nodes on one side of the bipartite network directly if they shared a node on the other side. By the end of the process, one side was eliminated. All network properties were calculated for the six projected networks (two projections for each of the predicted network, realised network without weights, and realised network with weights) as well. Structural erosion from the predicted to the realised network was tested using a paired Wilcoxon signed-rank test, comparing degrees of the same nodes across networks. To assess whether highly connected nodes were disproportionately affected, hub erosion was quantified as the Spearman rank correlation between predicted node degree and the number of links lost per node. Per ESP, the proportion of realised links classified as ES was calculated. Taxonomic differences in ES emphasis were tested using a Kruskal-Wallis rank-sum test, with post-hoc interpretation based on effect sizes rather than pairwise testing. The proportion of cross-type edges (ES–EDS) in the realised projection was compared to a null distribution generated by permuting ES/EDS labels across service nodes while preserving network topology through 10,000 permutations.

## Results

### Differences in predicted and realised knowledge networks

Scopus searches based on the predicted network resulted in 10,138 hits from 8,141 papers, of which 1,154 records from 882 publications were retained after LLM filtering. As a result, the predicted network, which comprised 47 non-isolated nodes connected by 103 links (Fig. 2A), decreased to 33 nodes and 58 links in the realised network (Fig. 3A), equalling 29.8% and 43.7% fewer nodes or links, respectively. ES nodes decreased from 18 to 10, EDS nodes from 9 to 4, and ESP nodes from 20 to 19 (Table 1). The EDS “Negative impact on beneficial organisms” exhibited the greatest loss between the predicted and realised networks, losing 100% of its 11 potential connections. Fourteen nodes were excluded from the realised network because they lost 100% of their connecting links: seven ES (e.g. cultural practices, climate regulation, fuel), four EDS (e.g. negative impacts on beneficial organisms, pest refuges, weeds), and one ESP (earthworms). Five ES/EDS nodes, such as pest control and pollination, did not lose any links (Supplementary tables 2-4).

**Figure 3.**
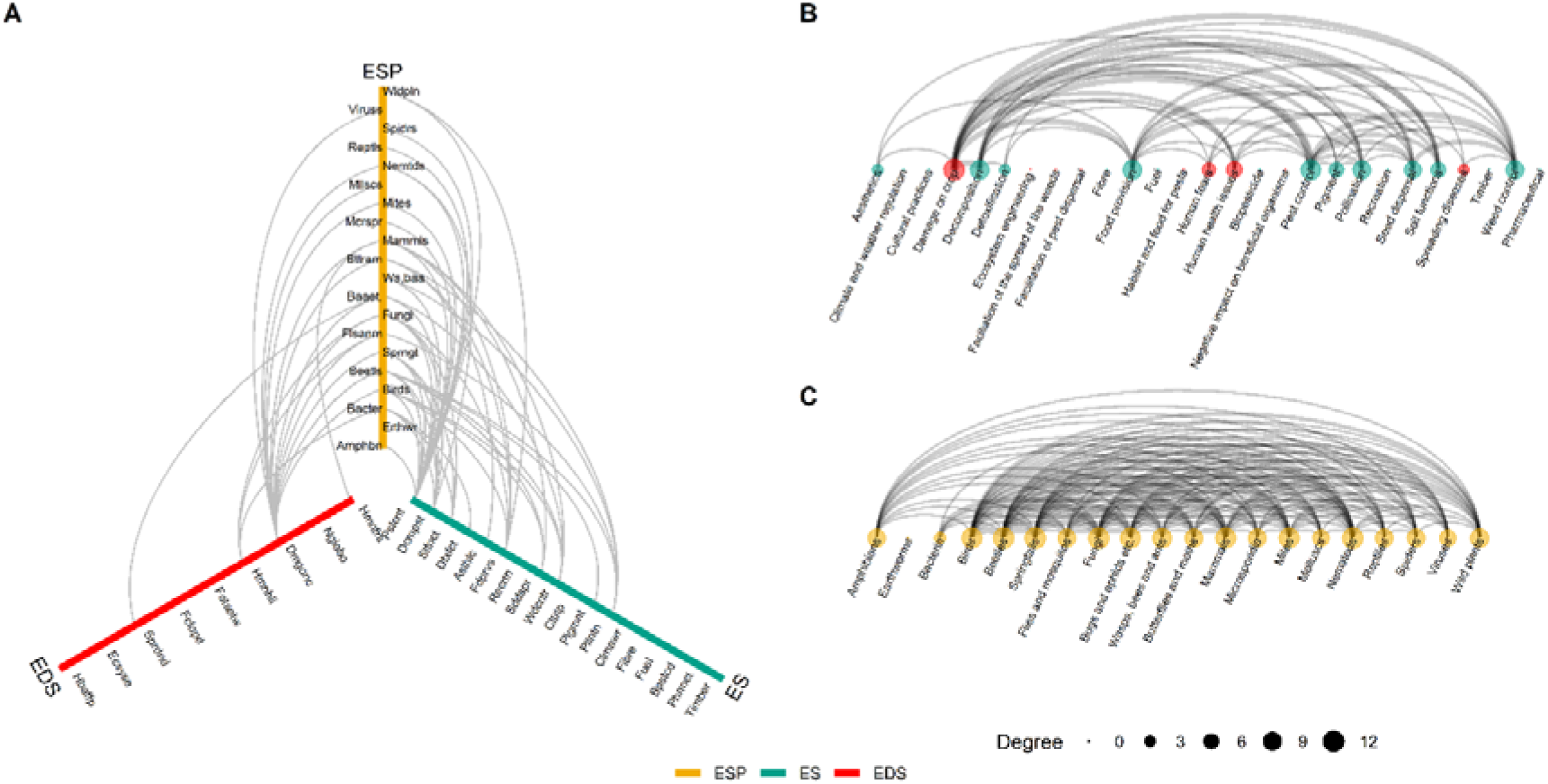
**(A)** Tripartite relationship between ecosystem services (ES), ecosystem disservices (EDS), and ecosystem service providers (ESP) in the realised network. Abbreviations are listed in Supplementary Table 1. **(B)** The first projection of the realised network connecting ES and EDS to each other, based on common links to ESP. Green dots represent ES, red dots EDS. Size of the dots is proportional to the degree (i.e. how many links it has to others) of a particular ES/EDS. **(C)** The second projection of the realised network connecting ESP to each other, based on common links to ES/EDS. Size of the dots are proportional to the degree (i.e. how many links it has to others) of a particular ESP.

**Table 1.** Summary of predicted and realised networks. Node counts refer to nodes with at least one link (non-isolated nodes). ES and EDS edges are counted in the main networks as links from ESPs to ES or EDS nodes, and in the ES/EDS projections as within-type (ES–ES or EDS–EDS) edges. ESP projection networks contain only ESP nodes.

| Network |  | Total nodes | ES nodes | EDS nodes | ESP nodes | Total edges | ES edges | EDS edges |
| --- | --- | --- | --- | --- | --- | --- | --- | --- |
| Predicted main network |  | 47 | 18 | 9 | 20 | 103 | 61 | 42 |
| Realised main network |  | 33 | 10 | 4 | 19 | 58 | 41 | 17 |
| Predicted projection | ES/EDS | 27 | 18 | 9 | 0 | 227 | 108 | 25 |
| Realised projection | ES/EDS | 14 | 10 | 4 | 0 | 51 | 27 | 3 |
| Predicted projection | ESP | 20 | 0 | 0 | 20 | 156 | NA | NA |
| Realised projection | ESP | 19 | 0 | 0 | 19 | 137 | NA | NA |

Despite a substantial loss of edges and nodes, the realised network remained structurally similar to the predicted one when judged at the adjacency level. The correlation between the upper triangles of the predicted and realised adjacency matrices was 0.73, and this similarity did not differ from a null model of random edge loss (random thinning; p = 1). Nonetheless, since most ES, EDS, and ESPs underwent a decrease in their degree from the predicted to the realised network, node-level connectivity was systematically reduced. Node degrees declined between predicted and realised networks (W = 666, p < 0.001), with degree loss positively associated with predicted degree (Spearman ρ = 0.27), indicating disproportionate erosion of hubs (Supplementary Figure 11).

Node-level loss of connectivity differed between ES and EDS; EDS nodes lost 69% of their predicted links on average, compared to 57.4% for ES nodes (Wilcoxon rank-sum test, W = 121, p = 0.020), with the mean loss of EDS being 2.78 links (SD = 3.19) and 1.11 links (SD = 0.96) for ES.

Generally, less support for the *a priori* predicted EDS links was found in the literature than for ES. Across all predicted links, ES were more likely to be realised in the literature than EDS, and, conversely, less likely to be lost when moving from the predicted to the realised network. The difference in realisation probability was modest but positive, ΔP(realised | ES – EDS) = 0.267 (95% CI 0.071–0.456), with the corresponding loss probabilities being 0.328 for ES links and 0.595 for EDS links. This asymmetry was strongly supported by label permutation tests (realisation: one□sided p = 0.006, two□sided p = 0.0091; loss: one□sided p = 0.0061) and Fisher’s exact test for loss (p = 0.009). The logistic regression of loss on link type yielded an odds ratio of 3.01 for EDS vs ES links (95% CI 1.35–6.93), indicating that predicted ES links were only about half as likely to be lost as EDS ones.

When the strength of evidence was considered by weighting links with the number of supporting studies, ES received substantially more attention than EDS. Across all predicted links, the total number of papers associated with ES links was 1003 compared with 151 for EDS links, corresponding to an ES:EDS evidence ratio of 6.64 (95% bootstrap CI 3.03–16.11), which significantly deviated from the expected 1. The zero-inflated negative binomial model indicated that, among realised links, ES links tended to have higher paper counts than EDS links (incidence rate ratio 3.31, 95% CI 1.35–8.09), while the zero-inflation component suggested non-significant differences in the probability that links are completely absent from the literature (odds ratio for ES vs EDS structural zero = 0.19, 95% CI 0–8.83).

The reduction in links between the predicted and realised networks also cascaded into the projected networks: the ES/EDS projection linking services and disservices through common ESPs declined from 227 to 51 edges between 27 and 14 nodes, respectively (Fig. 2B, Fig. 3B). Thus, link density declined substantially from the predicted to the realised knowledge network (from 0.65 to 0.15). In contrast, the second projection linking ESPs via shared ES or EDS remained highly connected and lost relatively few links, with declines from 156 to 137 edges between 20 and 19 nodes (respectively) (Fig. 2B, Fig. 3B).

### Inconsistent evaluation of ESPs

Most ESPs lost potential links in the realised network. Wild (non-crop, non-weed) plants, mammals, and birds lost 14, 8, and 6 of the original 16, 14, and 11 connections, respectively. Taxa tended to lose more EDS- than ES-related links from the predicted to the realised network, with an average decline of 1.00 ± 2.55 ES links and 1.25 ± 1.33 EDS links from the predicted to the realised network (Wilcoxon rank-sum test, W= 259, p = 0.057). Along with disproportionately losing links, the ES:EDS ratio of the nodes connected to ESPs increased in the realised networks, and these ratios differed significantly among ESPs (Kruskal–Wallis χ2 = 18, p = 0.456). This shift was particularly pronounced when links were weighted with the number of the published papers (Fig. 4C).

**Figure 4.**
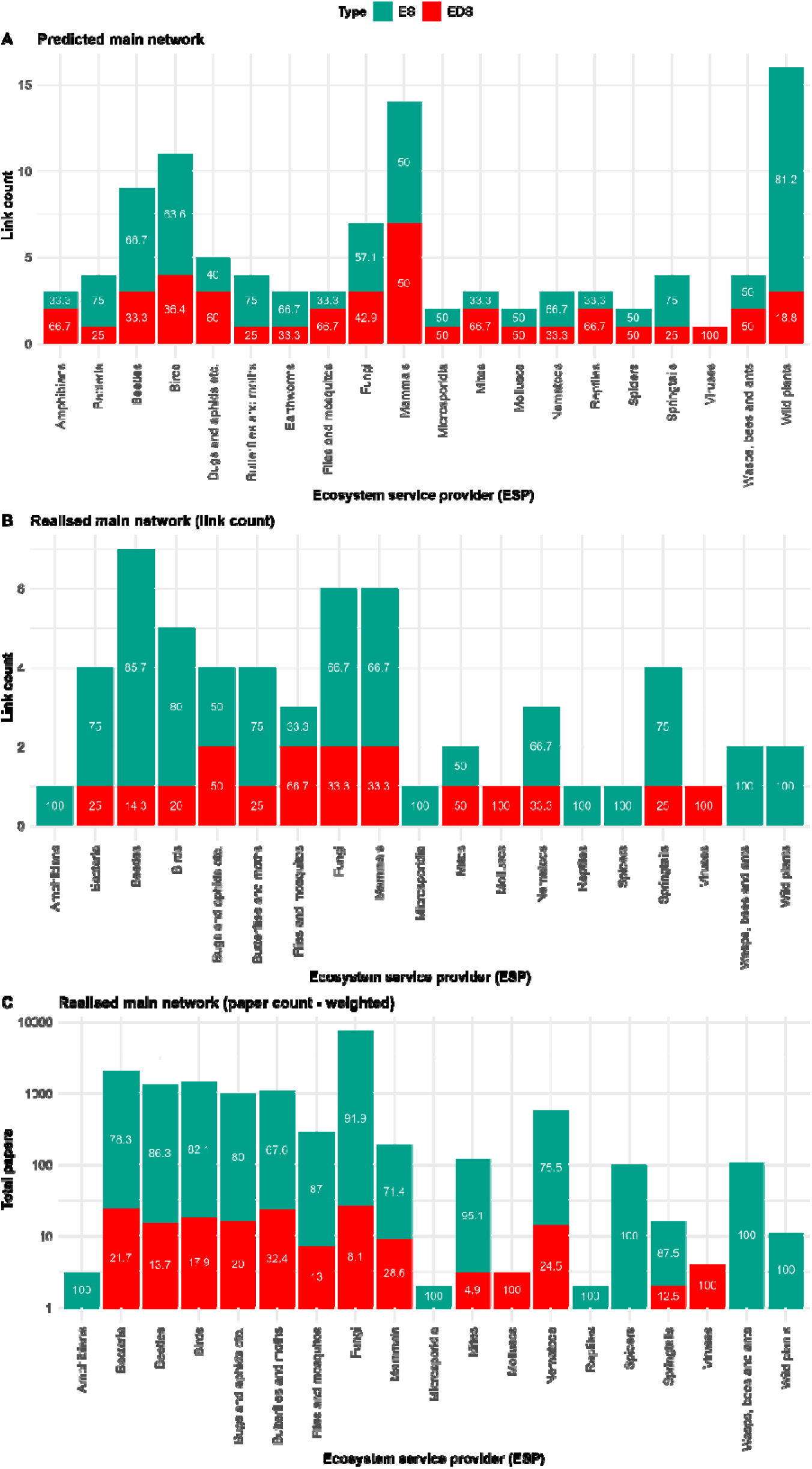
Distribution of ecosystem service (ES, green) and ecosystem disservice (EDS, red) links across ecosystem service providers (ESPs). **A**: predicted network showing conceptually plausible ES and EDS links. **B**: realised network showing the number of documented ES and EDS links per ESP (link count). **C**: realised network weighted by the total number of papers supporting each link type (paper count). Numbers within bars show the percentage of ES or EDS for each ESP. The shift toward higher ES proportions from predicted to realised networks, and the further amplification when weighted by evidence, indicates preferential documentation and replication of beneficial functions over disservices.

As a result, certain taxa became disproportionately framed as ES-only providers (Fig. 5). Yet, the realised network revealed strong variation among taxa in the proportion of their links classified as ecosystem services or disservices, with ES:EDS ratios ranging from 0 to 1. Importantly, 6 taxa were effectively framed as ES-only providers, having no realised EDS links despite non-zero ES link counts, i.e. wasps, bees, and ants (Hymenoptera), wild plants, amphibians, Microsporidia, reptiles, and spiders. Some taxa were excessively studied or understudied, compared to their predicted roles in crop systems. Weighting the realised network with the number of logged articles showed that most papers (310) dealt with fungi, of which 91.9% were in the context of ES. Although the absolute number of papers featuring Hymenoptera and spiders was lower (104 and 99, respectively), these two groups were considered to provide ES only. In contrast, only three papers discussed viruses and all were associated with EDS. Despite their clear role as ESPs and their high conservation value, the least studied organisms in relation to crop systems were amphibians and reptiles, documented in one and two papers, respectively (Fig. 5, Supplementary table 6). Some other relatively little-studied organisms, such as springtails, with potentially high ES values were also largely understudied.

**Figure 5.**
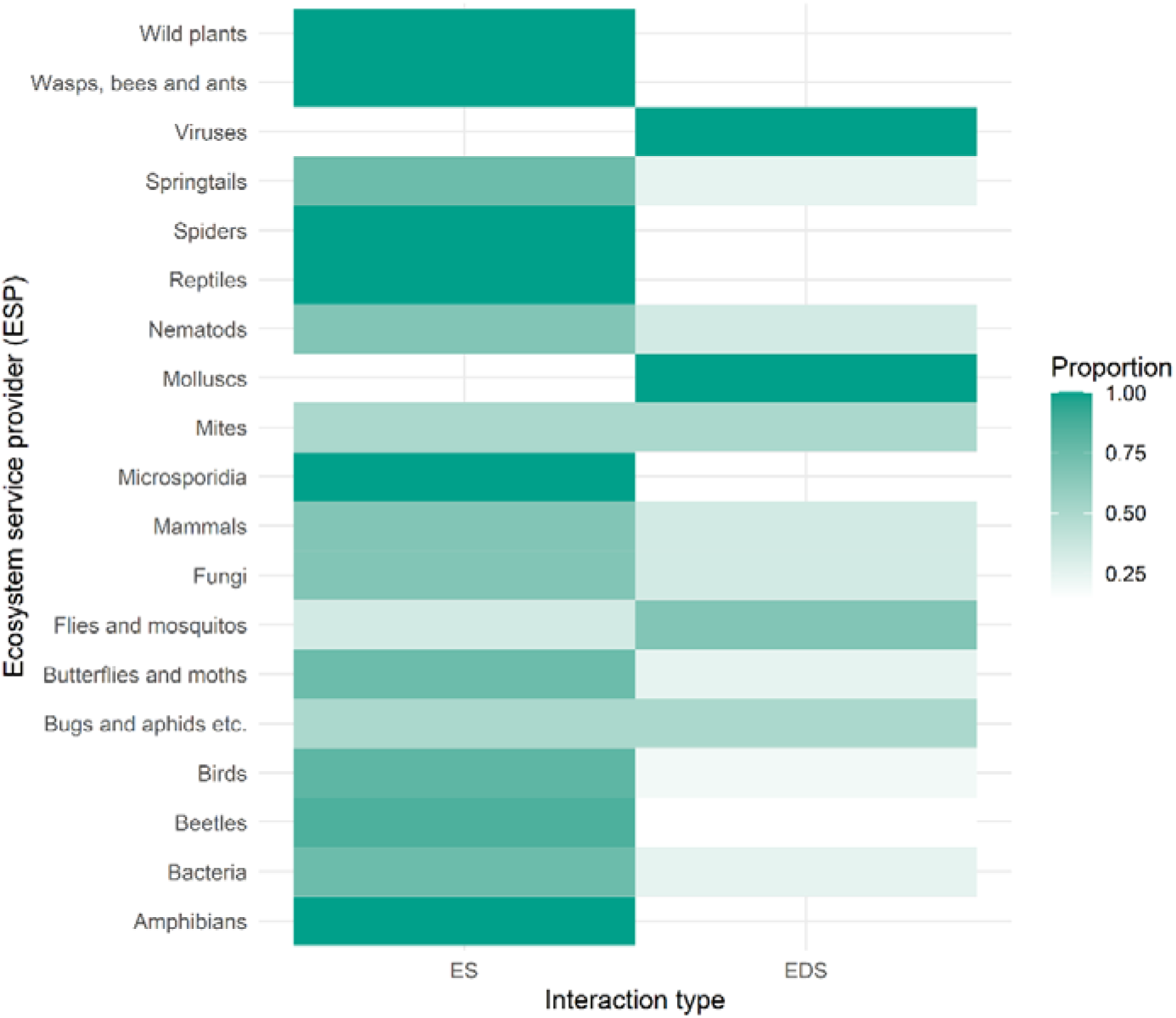
Taxon-specific linking to ES and ESDs. Colour depth indicates the proportion with which taxa are linked to ES or EDS.

### ES-EDS bundles through common ESPs

Projected side networks which link ES and EDS through common ESPs show bundled services and disservices, and are thus indicators of trade-offs (Fig. 3B, Fig. 4B, Supplementary Figures 13, 15, 17). The high link density of this projection of the predicted network suggests that most taxa are involved in delivering both ES and EDS. Yet, the link density declined substantially from the predicted to the realised knowledge network (from 0.65 to 0.15), mirroring the loss of edges in the main network (Table 1). Crucially, the proportion of cross-type ES–EDS edges in the realised projection was lower than expected under a null model that permuted ES/EDS labels among service nodes (observed proportion = 0.41, permutation test p = 0.266).

Out of all ES, pest control was linked to most EDS (4 in the realised knowledge network, through 11 ESPs), whilst decomposition was linked to the most ES (8 through 13 ESPs). As an EDS, “crop damage” had the greatest number of links to ES (9 in the realised knowledge network, through 30 ESPs), whilst it had 3 links (through 4 ESPs) to other EDS (Fig. 4B, Supplementary Table 5).

The other projected side networks that show which ESPs were linked through shared ES or EDS, indicates which ESPs are thought to be commonly involved in providing/causing the same ES/EDS (Fig. 3C, Fig. 4C, Supplementary Figures 14, 16). With a few exceptions, this projection of the predicted network was fully connected and, unlike in the projection between ES/EDS, few connections were lost (2.3 ± 1.91, mean ± SD) in the realised knowledge network (Supplementary Table 6). Already understudied taxa, such as earthworms and springtails, also lost a relatively high number of their links to other taxa.

## Discussion

Our structured analysis of knowledge networks reveals that ecosystem disservices are substantially underrepresented in the literature, which distorts the evidence base for cropland management and conservation planning. In addition to predicted ES connections being only about half as likely to disappear as EDS connections from the predicted to the realised knowledge networks, an even more striking pattern was the complete absence of information links (i.e. scientific papers) for certain disservice categories, such as negative impacts on beneficial organisms, pest refuge provision, and weed spread. Because of EDS links being disproportionately less supported by the literature, four of the nine predicted EDS nodes lost all connectivity, thus were not present in the realised network, compared to seven of the eighteen ES nodes.

Although our findings are consistent with earlier studies reporting disparities between ES and EDS in the scientific literature (Blanco et al., 2019), a more balanced pattern might nevertheless have been expected, given the substantial expansion of ES research over the past decade, particularly in Europe and the United States (McDonough et al., 2017; Urbina-Cardona et al., 2023), together with continued growth of the EDS literature (Guo et al., 2022). Yet, the fact that ES still received more than six times as many papers as disservices in our dataset than EDS suggests that cognitive biases, along with institutional filters, continue to shape the literature beyond simple publication bias. This is in line with Takacs and O’Brien (2023), who identified “hot” and “cold” topics in ecosystem service research, and suggests that similar filtering may also operate across disciplinary boundaries. Indeed, initial research emphasis may generate self-reinforcing visibility through citation networks, further concentrating attention on already established themes (Jeong et al., 2003; Merton, 1988). Besides these disciplinary and topical preferences, ecosystem services that are difficult to monetise, such as climate regulation and cultural practices, were more likely to be absent from our realised network, suggesting that services with clear economic value attract more research attention and funding (Nesbit et al., 2026). On the other hand, some ecosystem disservices can be conceptually ambiguous and therefore also difficult to quantify and align with agricultural productivity metrics driving funding priorities (Saunders, 2020), which may be one of the reasons they were underrepresented in our realised knowledge network.

Beyond service-disservice asymmetries, a taxonomic bias toward diurnal, conspicuous organisms and those of clear economic importance (Bentley, 1989; Wyckhuys et al., 2019b) emerged at the expense of the small-bodied, night-active and/or belowground fauna. In our case, mentions of cryptic soil fauna nearly vanished from the realised knowledge network, despite their central role in agroecosystem functioning (Wall et al., 2010; Wyckhuys et al., 2021); earthworms, important for weed seed burial, nutrient cycling, and soil structure maintenance, and indirectly for global food security (Fonte et al., 2023), were absent entirely, whereas springtails featured to minimal extent in the global ES literature. Amphibians illustrate this bias as well: although they remain marginal in our realised network, implying that they are seldom foregrounded in mainstream agricultural ecosystem service research. For instance, frogs, indeed, contribute to pest suppression in rice systems (Khatiwada et al., 2016; Zou et al., 2017). Most studies report that people are unaware of the value of organisms (Wyckhuys et al., 2019b, 2019a), which may also contribute to the systematic neglect of some taxa in research design and reporting. This bias may extend even to research practice, as biological control scientists tend to study day-active organisms even though nocturnal ones occasionally are functionally more important (Wyckhuys et al., 2024).

Whether taxa were more linked to ES or EDS was also imbalanced. Six taxa emerged as effectively ES-only providers despite conceptually plausible harmful roles. For instance, fungi were represented predominantly in an ES context in our realised network even though many fungi are major plant pathogens. This may point to a partial disciplinary or ideological disconnect between plant pathology and ecosystem service science, and more broadly to crop-protection traditions that do not readily frame taxa in terms of multifunctionality (Wyckhuys et al., 2023). Furthermore, spiders appeared in 99 papers but exclusively as pest control agents, with no documentation of the disservices they may cause, such as non-target or intraguild predation (Denno et al., 2004; Hodge, 1999), or generating human fears (e.g. Vance-Chalcraft et al., 2007). Similarly, some Hymenoptera, including sawflies and leaf-cutter bees, can also be plant-damaging in perennial or tree-based systems, although they were only considered as ES providers; mostly pollinators or biological control agents. Viruses illustrate a converse example: in our dataset they appeared only in relation to EDS, despite documented beneficial roles in biological control, including mycovirus-mediated suppression of forest diseases (Muñoz-Adalia et al., 2016). Our results thus demonstrate that beneficial functions of organisms are widely recognised but are not always evaluated together with possible costs, trade-offs, or other decision-relevant endpoints, which, in turn, is likely to limit a full understanding of net outcomes in practice (Kleijn et al., 2019; Naranjo et al., 2015).

Our analysis of network projections demonstrated contrasting patterns that clarify how this taxonomic bias operates. The ESP projection, linking taxa through shared services and disservices, retained 88% of edges, confirming that researchers recognise organisms’ multifunctionality. However, the ES/EDS projection, directly linking services and disservices through common providers, collapsed dramatically, losing 78% of edges. This asymmetry indicates that whilst multifunctionality is acknowledged at the organism level, the co-occurrence of services and disservices through the same organisms is rarely studied. Indeed, cross-type ES–EDS edges were underrepresented relative to random expectation, demonstrating a systematic, although most likely unconscious, avoidance of trade-off research.

The reasons for this taxonomic bias are likely to be multifaceted. Since most people value the presence of most organisms negatively (Gurung, 2003; Lyytimäki, 2014; McLellan and Shackleton, 2019, but see Martin and Doucet, 2022), the perceived importance of biodiversity conservation could be further weakened by highlighting EDS (Shapiro and Báldi, 2014; Villa et al., 2014). Thus, the service-focused research may reflect academic and institutional priorities to (over)compensate processes in common media, rather than farmer information needs. Confirmation bias (Kahneman et al., 2011) likely reinforces these patterns, as researchers working on conservation biological control or habitat enhancement may unconsciously seek evidence supporting beneficial effects whilst overlooking potential harms. Similarly, optimistic bias, where one selectively integrates positive information while discounting negative evidence during belief updating, may also be influential (Kozlov et al., 2014). Whilst these taxonomic biases in ecosystem service research do exist both in terrestrial (Noriega et al., 2018) and aquatic environments (Nabout et al., 2023), this selective framing means practitioners lack information about context-dependencies determining net outcomes of habitat enhancement strategies (Tschumi et al., 2018).

### Implications for adaptive management

To unlock agroecological transitions in global agri-food systems and ecologically intensify agriculture, a comprehensive understanding of interactions across full taxonomic communities is imperative (Erktan et al., 2024; Tscharntke et al., 2012; Wyckhuys et al., 2025). The knowledge gaps we unveiled here pose practical limitations for management aiming to optimise multiple ecosystem functions simultaneously (Mkenda et al., 2019). Importantly, without systematic disservice quantification, scientists and farmers alike are unable to anticipate net outcomes across contexts, and the predominantly optimistic framing creates risks when translated to management. The resulting risk aversion and impatience hamper farmers’ adoption of sustainable farming tactics (Simutowe et al., 2024). If researchers selectively document benefits whilst underinvestigating risks, they provide inadequate information for risk-averse farmers facing adoption decisions. If interventions assume purely beneficial outcomes, unanticipated disservices may require corrections negatively affecting farmers’ welfare and livelihood security (Herd-Hoare and Shackleton, 2020). At the same time, decreasing this bias may further promote the use of the ES/EDS framework in decision-making (Laurans et al., 2013).

The identified gaps have direct implications for sustainable farming tactics such as agroecology and biological control, i.e. foundational elements of integrated pest management (IPM) (Deguine et al., 2021). For example, pest control appeared in 410 papers across 14 connections, making it the most studied ES-ESP link in our realised network, but it was linked to only three EDS through 10 ESPs. Thus, these interactions rarely receive joint investigation despite obvious relevance for optimising biological control, which may obscure critical trade-offs. Yet, the two most important cropland ES and EDS, pest control by natural enemies and crop damage by pests were intertwined, indicating a difficult path for plant protection and ecological engineering projects (Gurr et al., 2004) that aim to maximise biological pest control without increasing crop damage. Promoting taxa that are linked to a low number of EDS (e.g. spiders) and which are bundled with other taxa with a high ES:EDS ratio (e.g. non-crop plants) can resolve this (Kremen and Miles, 2012) as long as the strengths of ES and EDS are the same.

Similarly, we predicted “wild plants” to be the most connected ESP (degree = 16), with 13 links to ES. Thus, by providing nutrition or shelter to natural enemies, wild plants were the most beneficial organisms in crop systems. In the realised network, wild plants were related only to pest control, food provision, and pollination ES, indicating a unilateral representation of research on the benefits of flower strips, field margins, and cover crops in conservation biological control. Indeed, most studies in this area focus on benefits (Marshall and Moonen, 2002; Kovács-Hostyánszki et al., 2016), while potential harms, such as wildflower margins facilitating weed spread or disrupting pest control (e.g. Genty et al., 2026), are rarely examined (Ganser et al., 2019; Gontijo, 2019), which thereby hampers the development of successful IPM programs. Similarly, although there is evidence that flower strips can harbour alternative prey or competing predators, potentially reducing biological control effectiveness (e.g. Martin et al., 2013), papers investigating organisms’ negative impacts on other, beneficial, organisms were not present in our realised network. On the other hand, weed ecology may have a separate literature pool that remained untapped since we did not use the term “weed” in our searches.

### Research priorities and institutional reform

Addressing EDS alongside ES is essential for reducing their negative effects while maintaining ecosystem resilience (Lyytimäki, 2015) and informing sustainability-oriented decision-making (Schaubroeck, 2017, Shackleton et al., 2016). To compensate systematic biases requires institutional changes (Vanloqueren and Baret, 2017): publication and funding systems must evolve to valorise comprehensive assessments that document both services and disservices, null results, and negative findings. Although existing conservation frameworks are biased towards positive values, plural valuation frameworks are needed that account for both benefits and harms (Oostvogels et al., 2024). This can be tackled through mandatory reporting requirements to address potential biases (Zvereva and Kozlov, 2021), involving requirements to explicitly address potential disservices, report effect sizes and uncertainty for both positive and negative outcomes, and avoid selective reporting of favourable results. Furthermore, funding agencies could also require explicit disservice investigation as a condition of service-focused grants.

Integration of traditional and local knowledge with scientific research could help identify context-dependent factors that determine when organisms shift from net service to disservice provision. Insights are also needed into the practical implications of these disservices and their cascading impacts on income, farm revenue, or return on investment (Kleijn et al., 2019). Yet, such knowledge integration remains limited. This gap becomes particularly important given systematic disagreement between farmers and researchers about which adoption obstacles matter most, suggesting service-focused research may reflect academic priorities rather than farmer information needs (Chaplin-Kramer et al., 2019; Geertsema et al., 2016; Maas et al., 2021). In any case, involving farmers in identifying which disservices matter in their specific contexts could help correct mismatches between research priorities and farmer information needs, leading to biodiversity-based management strategies that better reflect locally relevant risks, adoption barriers, and economic trade-offs (Velado-Alonso et al., 2024).

Our work helps to set the following research priorities: (1) to operationalize ecological intensification (Tittonell, 2014; Vanbergen et al., 2020), the missing disservice categories need to be systematically uncovered and investigated; (2) given their documented ecological importance, severely understudied taxa, earthworms, amphibians, reptiles, and springtails, need a comprehensive ES/EDS assessment in a complex on-farm biostructure framework (Wyckhuys et al., 2025); (3) trade-off quantification addressing the collapsed ES/EDS projection should become standard practice; (4) context-dependency research explaining when organisms provide net benefits versus costs across varying management and environmental conditions. Network models, similar to ours, could be used to assess multiple ES and EDS by building conceptual maps (García-Díaz et al., 2025), comparing them to evidence-based knowledge networks, and predicting their environmental and socio-economic outcomes.

Some limitations should be considered when interpreting our results. First, the keyword co-occurrence approach cannot fully disentangle true knowledge gaps from publication bias, nor does it capture the quantitative effect sizes that are often required for direct management application. Although the complete disappearance of certain disservice categories and the observed six-fold imbalance in evidence are unlikely to be attributable to methodological artefacts alone, the keywords employed were necessarily selective and therefore unavoidably arbitrary. In addition, the exclusion of pesticide efficacy papers may have disproportionately removed studies that framed pest problems and related harms as disservices, even when these were introduced as the rationale for control, thereby further reducing the visibility of some EDS links in the realised network. Furthermore, even though our machine learning approach exhibited high levels of human agreement, it could have led to the accidental omission of relevant studies. Meanwhile, manual screening itself is not free of errors (Gartlehner et al., 2020) and exhibited 5-7% disagreement among human assessors.

### Conclusions

Our work unveils a substantial bias toward ecosystem services in agroecosystem research, with disservice links experiencing 1.8-fold higher loss probability and 6.6-fold lower evidence accumulation. While taxa are recognised as multifunctional, certain groups remain systematically underrepresented, and the specific bundling of services with disservices is likewise overlooked. This biased evidence base can limit the success of ecological intensification by creating unrealistic expectations about intervention outcomes and reducing adaptive capacity across heterogeneous production contexts. Effective landscape management and agricultural interventions therefore require frameworks that evaluate trade-offs more holistically and clarify how interacting ecological processes, rather than single services or disservices in isolation, shape yields and broader social-ecological outcomes in crop fields. Addressing ecosystem disservices alongside ecosystem services is essential for maintaining ecosystem resilience and supporting more context-sensitive decision-making, but doing so will also require socio-institutional reform, including updated funding priorities, publication standards, methodological protocols, and stakeholder engagement

### Declaration of AI Use

During the preparation of this work, the authors used a locally deployed Large Language Model (Qwen2.5-14B) to automate literature screening, having first validated its accuracy against a human-curated dataset. The process is detailed in the Materials and Methods section of the manuscript. Additionally, AI tools were used to rephrase text for readability; the authors reviewed all AI-assisted writing and take full responsibility for the manuscript’s final content.

## Acknowledgements

This work was supported by a “111 Program” grant to Minsheng You and Geoff Gurr at Fujian Agriculture & Forestry University (FAFU). Gabor Pozsgai was supported by a postdoctoral fellowship by the State Key Laboratory of Agriculture and Forestry Biosecurity at Fujian Agriculture and Forestry University in China at FAFU. Liette Vasseur supported HV and MH through her UNESCO Chair on Community Sustainability.

## Author Contributions

The idea was conceived by G.P., G.L., L.V., G.M.G., M.G., O.R., K.W., S.W., and M.Y. Data collection procedure, statistical analysis, and data representation were designed and conducted by G.P. and I.B.F. Each contributor participated in data collection. First draft of the text was written by G.P. and K.W., with figures drawn by I.B.F, then edited by G.L., M.Y., G.G., O.R, L.V., M.G., and S.W. The final version of the manuscript was edited by G.P., K.W, I.B.F., G.L., L.V., G.G. and M.Y. All contributors participated in the filtering process and read the final version of the manuscript.

## Data and code availability

All raw data and R code used for data manipulation and network generation will be made available on a public repository (https://github.com/pozsgaig/cognitive_bias) upon acceptance.

