## Supplementary material for "Ecosystem disservices are underrepresented in literature on annual crop agroecosystems": Supplementary_material.html

#### 2026-06-07

- Hypothetical network input
- AI Model Performance (LLMs)
- ML Model Performance Comparison (Non-LLMs)
- Node degree loss
- Network projections

Gábor Pozsgai^1,2,3\*§^, Kris A.G. Wyckhuys^2,3,4,5,6§^, Ibtissem Ben Fekih^3,7,8§^, Liette Vasseur^2,3,9^, Geoff Gurr^2,3,10^, Mark Goettel^2,3,11^, Heather VanVolkenburg^9^, Sunita Pandey^12^, Jamile Queiroz-Sousa^13^, Pingyang Zhu^14^, Mohammed Abul Monjur Khan^15^, Titus Imboma^2,3,16^, Margaret Hughes^17,18^, Jian Liu1^9^, Syed Rizvi^20^, Olivia Reynolds^2,3,10^, Hafiz Sohaib Ahmed Saqib^2,3,21^, Stephen Wratten^2,3,22†^, Jie Zhang^2,3,23^, Wenwu Zhou^24^, Francisco Javier Garcia^25^, Lucas Alexander Shuttleworth^26^, Li-Lin Chen^2,3^, Gábor L. Lövei^2,3,27\*^, Minsheng You^2,3\*^

^1^Centre for Applied Economics of the Atlantic, University of Azores, Angra do Heroismo, Portugal

^2^Joint International Research Laboratory of Ecological Pest Control, Ministry of Education, Fuzhou 350002, China

^3^State Key Laboratory of Agriculture and Forestry Biosecurity, Institute of Applied Ecology, Fujian Agriculture and Forestry University, Fuzhou 350002, China

^4^Chrysalis Consulting, Danang, Vietnam

^5^School of the Environment, The University of Queensland, Saint Lucia, Australia

^6^Institute of Plant Protection, Chinese Academy of Agricultural Sciences (CAAS), Beijing, China

^7^Institute of Environmental Microbiology, College of Resources and Environment, Fujian Agriculture and Forestry University, Fuzhou, China

^8^Functional and Evolutionary Entomology, Terra, Gembloux Agro-Bio Tech, University of Liege, Passage des Déportés 2, 5030 Gembloux, Belgium.

^9^UNESCO Chair on Community Sustainability: From Local to Global, Dept. Biol. Sci., Brock University, Canada

^10^Gulbali Institute, School of Agriculture, Environment and Veterinary Sciences, Charles Sturt University, Orange NSW 2800, Australia

^11^Agriculture and Agri-Food Canada, Lethbridge Research Centre, Lethbridge, AB, Canada

^12^Ministry of Agriculture and Livestock Development, Singhadurbar, Nepal

^13^Instituto de Biociências de Botucatu (IBB), Universidade Estadual Paulista Júlio de Mesquita Filho (UNESP), Botucatu, São Paulo, Brazil

^14^College of Life Sciences, Zhejiang Normal University, Jinhua 321004, China

^15^Department of Entomology, Faculty of Agriculture, Bangladesh Agricultural University, Mymensingh-2202, Bangladesh

^16^Ornithology Section of the Zoology Department, National Museums of Kenya, PO Box 40658- 00100, GPO. Nairobi, Kenya
^17^Department of Biological Sciences, University of Calgary, Calgary, AB, Canada

^18^Department of Biological Sciences, Brock University, St. Catharines, ON, Canada

^19^Artificial Intelligence and Cyber Futures Centre, Charles Sturt University, Orange NSW 2800, Australia

^20^Applied Biosciences, Macquarie University, Sydney, Australia

^21^Yunnan Key Laboratory of Forest Ecosystem Stability and Global Change, Xishuangbanna Tropical Botanical Garden, Chinese Academy of Sciences, Menglun, Mengla, Yunnan 666303, China

^22^Bio-Protection Research Centre, Lincoln University, New Zealand

^23^Fujian Key Laboratory for Monitoring and Integrated Management of Crop Pests, Fujian Engineering Research Center for Green Pest Management, Institute of Plant Protection, Fujian Academy of Agricultural Sciences, Fuzhou 350013, China

^24^State Key Laboratory of Rice Biology, Key Laboratory of Molecular Biology of Crop Pathogens and Insects, Ministry of Agriculture, Zhejiang University, Hangzhou, China

^25^Independent researcher, Spain

^26^National Institute of Agricultural Botany (NIAB), Department of Pest & Pathogen Ecology, East Malling, Kent ME19 6BJ United Kingdom

^27^Department of Agroecology, Flakkebjerg Research Centre, Aarhus University, 4200 Slagelse, Denmark

^§^These authors contributed equally to this work.

^†^Deceased; contributed to this work prior to passing

### Hypothetical network input

Table 1: Contents of the hypothetical network input workbook (prednet.xlsx).

| ESP\_name | ESP | ES\_ESD\_name | Type | Keyword | Rationale/Example | Comments |
| --- | --- | --- | --- | --- | --- | --- |
| Earthworms | Annelida | Damage on crop | EDS | Crop damage | Earthworms can mechanically damage crop roots | NA |
| Birds | Bird\* | Damage on crop | EDS | Crop damage | Birds eat crop seeds | NA |
| Beetles | Coleoptera | Damage on crop | EDS | Crop damage | Herbivorous beetles feeding on crop species. | NA |
| Springtails | Collembola | Damage on crop | EDS | Crop damage | Garden springtail (Bourltiella hortensis) can feed on crop leaves. | NA |
| Flies and mosquitos | Diptera | Damage on crop | EDS | Crop damage | Cabbage fly eating Brassicaceae species. | NA |
| Fungi | Fung\* | Damage on crop | EDS | Crop damage | Claviceps species infecting wheat. | NA |
| Bugs and aphids etc. | Hemiptera | Damage on crop | EDS | Crop damage | Aphids sucking on crop. | NA |
| Butterflies and moths | Lepidoptera | Damage on crop | EDS | Crop damage | Caterpillars chewing on crop. | NA |
| Mammals | Mammal\* | Damage on crop | EDS | Crop damage | Voles eating crop seeds. | NA |
| Mites | Mite\* | Damage on crop | EDS | Crop damage | Mites causing direct damage on crop | NA |
| Molluscs | Mollusc\* | Damage on crop | EDS | Crop damage | Slugs eating crop leaves | NA |
| Nematods | Nematod\* | Damage on crop | EDS | Crop damage | Nematode damage on potatoes. | NA |
| Viruses | Virus\* | Damage on crop | EDS | Crop damage | Virus infections on tobacco | NA |
| Wild plants | Wild plant\* | Damage on crop | EDS | Crop damage | Competition with crops | NA |
| Bugs and aphids etc. | Hemiptera | Spreading diseases | EDS | Disease vector | May be viros vectors for crops | NA |
| Wasps, bees and ants | Hymenoptera | Ecosystem engineering | EDS | Ecosystem engineering | Ants move soil | NA |
| Mammals | Mammal\* | Ecosystem engineering | EDS | Ecosystem engineering | Moles dig holes | NA |
| Amphibians | Amphibian\* | Human fears | EDS | Fear\* | Frogs may induce disgust in humans. | NA |
| Mammals | Mammal\* | Human fears | EDS | Fear\* | Fear of mice or large mammals | NA |
| Reptiles | Reptile\* | Human fears | EDS | Fear\* | Disgust of snakes. | NA |
| Bacteria | Bacteria | Human health issues | EDS | Human health | Bacteria causing diseases. | NA |
| Flies and mosquitos | Diptera | Human health issues | EDS | Human health | Tabanidae species spread human disaeses from livestock | NA |
| Fungi | Fung\* | Human health issues | EDS | Human health | Claviceps species are poisonous to humans. | NA |
| Mammals | Mammal\* | Human health issues | EDS | Human health | Badgers spreading bovine TBC. | NA |
| Wild plants | Wild plant\* | Human health issues | EDS | Human health | Hayfiever | NA |
| Birds | Bird\* | Facilitation of pest dispersal | EDS | Pest dispersal | Foretic pests | NA |
| Mammals | Mammal\* | Facilitation of pest dispersal | EDS | Pest dispersal | Foretic pests | NA |
| Wild plants | Wild plant\* | Habitat and food for pests | EDS | Pest refuge | Wild plants provide refuge for pests | NA |
| Amphibians | Amphibian\* | Negative impact on beneficial organisms | EDS | Reduc\* benefit\* | Frogs may predate on parasites or predators of pests. | NA |
| Birds | Bird\* | Negative impact on beneficial organisms | EDS | Reduc\* benefit\* | Birds predating on parasites or predators of pests. | NA |
| Beetles | Coleoptera | Negative impact on beneficial organisms | EDS | Reduc\* benefit\* | Predatory beetles eat other predatory beetles | NA |
| Fungi | Fung\* | Negative impact on beneficial organisms | EDS | Reduc\* benefit\* | Entomopathogenic fungi infecting predatory beetle species | NA |
| Bugs and aphids etc. | Hemiptera | Negative impact on beneficial organisms | EDS | Reduc\* benefit\* | Predatory Hemiptera preying on other predatory insect species. | NA |
| Wasps, bees and ants | Hymenoptera | Negative impact on beneficial organisms | EDS | Reduc\* benefit\* | Parasitic wasp parasitizing predatory insect species. | NA |
| Mammals | Mammal\* | Negative impact on beneficial organisms | EDS | Reduc\* benefit\* | Predatory mammals consuming predatory insects. | NA |
| Microsporidia | Microsporidia | Negative impact on beneficial organisms | EDS | Reduc\* benefit\* | Microsporidia can infect bees and other beneficial insects | NA |
| Mites | Mite\* | Negative impact on beneficial organisms | EDS | Reduc\* benefit\* | Predatory mites may consume other predators. | NA |
| Reptiles | Reptile\* | Negative impact on beneficial organisms | EDS | Reduc\* benefit\* | Lizards predating on predatory insects | NA |
| Spiders | Spider\* | Negative impact on beneficial organisms | EDS | Reduc\* benefit\* | Intraguild predation | NA |
| Birds | Bird\* | Faciliation of the spread of the weeds | EDS | Spread\* weed\* | Seeds of weed may stick to the legs or feathers of birds that later visit cropfields. | NA |
| Beetles | Coleoptera | Faciliation of the spread of the weeds | EDS | Spread\* weed\* | May move seeds of weed to cropfields. | NA |
| Mammals | Mammal\* | Faciliation of the spread of the weeds | EDS | Spread\* weed\* | Weed seeds can stuck in mammals fur. | NA |
| Birds | Bird\* | Aesthetics | ES | Aesthetic\* | Colourful birds and birdsongs | NA |
| Butterflies and moths | Lepidoptera | Aesthetics | ES | Aesthetic\* | Colourful butterflies | NA |
| Mammals | Mammal\* | Aesthetics | ES | Aesthetic\* | Charismatic mammals | NA |
| Wild plants | Wild plant\* | Aesthetics | ES | Aesthetic\* | Colourful flowers | NA |
| Wild plants | Wild plant\* | Medicine/organic pesticide resources | ES | Biopesticide | Nettle used as organic pesticide | NA |
| Wild plants | Wild plant\* | Climate and weather regulation | ES | Climate regulation | Wild plants provide shaded, moist microclimate | NA |
| Birds | Bird\* | Cultural practices | ES | Cultural practices | Birdwatching | NA |
| Mammals | Mammal\* | Cultural practices | ES | Cultural practices | Wildlife photography | NA |
| Wild plants | Wild plant\* | Cultural practices | ES | Cultural practices | Tourists visit sunflower fields | NA |
| Earthworms | Annelida | Decomposition | ES | Decomposition | Earthworms eating detritus in soil | NA |
| Bacteria | Bacteria | Decomposition | ES | Decomposition | Bacteria decompose organic material in soil | NA |
| Beetles | Coleoptera | Decomposition | ES | Decomposition | Cadaver and dung beetles | NA |
| Springtails | Collembola | Decomposition | ES | Decomposition | Springtails process fallen leaves | NA |
| Fungi | Fung\* | Decomposition | ES | Decomposition | Fungi decompose dead plant material | NA |
| Nematods | Nematod\* | Decomposition | ES | Decomposition | Mechanical processing of dead plant material | NA |
| Bacteria | Bacteria | Detoxification | ES | Detoxification | Bacteria break down toxins in soil | NA |
| Wild plants | Wild plant\* | Detoxification | ES | Detoxification | Brachiaria species detoxify aluminium polluted soil | NA |
| Mammals | Mammal\* | Ecosystem engineering | ES | Ecosystem engineering | Is there any benefit in cropfields? | NA |
| Wild plants | Wild plant\* | Fibre | ES | Fibre\* | Cover crops harvested for fibre | NA |
| Birds | Bird\* | Food provision | ES | Food | Gamebirds | NA |
| Fungi | Fung\* | Food provision | ES | Food | Mushroom picking | NA |
| Butterflies and moths | Lepidoptera | Food provision | ES | Food | Some larger caterpillars are eaten | NA |
| Mammals | Mammal\* | Food provision | ES | Food | Hunting in cropfields | NA |
| Molluscs | Mollusc\* | Food provision | ES | Food | Edible snails | NA |
| Wild plants | Wild plant\* | Food provision | ES | Food | Flax for oil | NA |
| Wild plants | Wild plant\* | Fuel | ES | Fuel | Cover crops grown for biofuel | NA |
| Amphibians | Amphibian\* | Pest control | ES | Pest control | Frogs predating on pests. | NA |
| Birds | Bird\* | Pest control | ES | Pest control | Birds predate on caterpillars | NA |
| Beetles | Coleoptera | Pest control | ES | Pest control | Ground beetles and ladybirds aphid predation | NA |
| Springtails | Collembola | Pest control | ES | Pest control | Collembola consume fungal spores | NA |
| Fungi | Fung\* | Pest control | ES | Pest control | Entomopathogenic fungi | NA |
| Bugs and aphids etc. | Hemiptera | Pest control | ES | Pest control | Assasin bugs preadte on pests | NA |
| Wasps, bees and ants | Hymenoptera | Pest control | ES | Pest control | Parasitoid wasps | NA |
| Mammals | Mammal\* | Pest control | ES | Pest control | Moles eating grubs | NA |
| Microsporidia | Microsporidia | Pest control | ES | Pest control | Entomopathogenic species | NA |
| Mites | Mite\* | Pest control | ES | Pest control | Predatory mites may consume pests. | NA |
| Nematods | Nematod\* | Pest control | ES | Pest control | Predatory nematods consuming pest nematods | NA |
| Reptiles | Reptile\* | Pest control | ES | Pest control | Lizards predating on pests | NA |
| Spiders | Spider\* | Pest control | ES | Pest control | Spiders predating on pests | NA |
| Wild plants | Wild plant\* | Pest control | ES | Pest control | Secondary volatiles repel pests | NA |
| Wild plants | Wild plant\* | Medicine/organic pesticide resources | ES | Pharmaceutical | Salix species for Aspirin | NA |
| Beetles | Coleoptera | Pigment | ES | Pigment\* | Herbs or medical plants as cover crops | NA |
| Wild plants | Wild plant\* | Pigment | ES | Pigment\* | Dye made of scarlet beetles | NA |
| Beetles | Coleoptera | Pollination | ES | Pollination | Melighetes species pollinate oilseed rape | NA |
| Flies and mosquitos | Diptera | Pollination | ES | Pollination | Hoverflies pollinate plants | NA |
| Wasps, bees and ants | Hymenoptera | Pollination | ES | Pollination | Bees and bumblebees pollinate plants | NA |
| Butterflies and moths | Lepidoptera | Pollination | ES | Pollination | Moths pollinate some plants | NA |
| Birds | Bird\* | Recreation | ES | Recreation\* | Birdwatching | NA |
| Mammals | Mammal\* | Recreation | ES | Recreation\* | Wildlife photography | NA |
| Wild plants | Wild plant\* | Recreation | ES | Recreation\* | Tourism | NA |
| Birds | Bird\* | Seed dispersal | ES | Seed dispersal | General seed dispersal, excluding weed species | NOT weed |
| Beetles | Coleoptera | Seed dispersal | ES | Seed dispersal | Seed eating carabidae disperse cover crop seeds | NOT weed |
| Mammals | Mammal\* | Seed dispersal | ES | Seed dispersal | Mammals eating fruits disperse cover crop seeds | NOT weed |
| Earthworms | Annelida | Soil functions | ES | Soil function\* | Earthworms eating detritus in soil. | NA |
| Bacteria | Bacteria | Soil functions | ES | Soil function\* | Nitrogen fixation | NA |
| Springtails | Collembola | Soil functions | ES | Soil function\* | Collembola aerating soil | NA |
| Fungi | Fung\* | Soil functions | ES | Soil function\* | Soil fungi improve water regime | NA |
| Wild plants | Wild plant\* | Timber | ES | Timber | Trees providing timber in agroforestry | NA |
| Birds | Bird\* | Weed control | ES | Weed control | Weed seed predation | NA |
| Beetles | Coleoptera | Weed control | ES | Weed control | Weed seed predation | NA |
| Bugs and aphids etc. | Hemiptera | Weed control | ES | Weed control | Weed seed predation | NA |
| Mammals | Mammal\* | Weed control | ES | Weed control | Weed seed predation | NA |

### AI Model Performance (LLMs)

Figure 1: Metrics for AI: Claude

Figure 2: Metrics for AI: GPT\_5-2

Figure 3: Metrics for AI: Gemini\_3

Figure 4: Metrics for AI: Ollama

Figure 5: Metrics for AI: Sonar

### ML Model Performance Comparison (Non-LLMs)

Figure 6: Metrics for Logistic

Figure 7: Metrics for RandomForest

Figure 8: Metrics for XGBoost

Figure 9: Metrics for NaiveBayes

Figure 10: Metrics for SVM

### Node degree loss

Table 2: ES nodes: degree in predicted and realised networks and total realised papers per node.

| Node | Abbr. | Pred. degree | Real. degree | Real. total papers |
| --- | --- | --- | --- | --- |
| Aesthetics | Asthtc | 4 | 1 | 1 |
| Biopesticide | Bpstcd | 1 | 0 | 0 |
| Climate and weather regulation | Clmawr | 1 | 0 | 0 |
| Cultural practices | Cltrlp | 3 | 0 | 0 |
| Decomposition | Dcmpst | 6 | 5 | 130 |
| Detoxification | Dtxfct | 2 | 1 | 8 |
| Fibre | Fibre | 1 | 0 | 0 |
| Food provision | Fdprvs | 6 | 5 | 200 |
| Fuel | Fuel | 1 | 0 | 0 |
| Pest control | Pstcnt | 14 | 14 | 410 |
| Pharmaceutical | Phrmct | 1 | 0 | 0 |
| Pigment | Pigmnt | 2 | 1 | 1 |
| Pollination | Pllntn | 4 | 4 | 154 |
| Recreation | Recrtn | 3 | 0 | 0 |
| Seed dispersal | Sddspr | 3 | 3 | 10 |
| Soil functions | Slfnct | 4 | 3 | 74 |
| Timber | Timber | 1 | 0 | 0 |
| Weed control | Wdcntr | 4 | 4 | 15 |

Table 3: EDS nodes: degree in predicted and realised networks and total realised papers per node.

| Node | Abbr. | Pred. degree | Real. degree | Real. total papers |
| --- | --- | --- | --- | --- |
| Damage on crop | Dmgonc | 14 | 12 | 106 |
| Ecosystem engineering | Ecsyse | 2 | 0 | 0 |
| Faciliation of the spread of the weeds | Fotsotw | 3 | 0 | 0 |
| Facilitation of pest dispersal | Fclopd | 2 | 0 | 0 |
| Habitat and food for pests | Hbaffp | 1 | 0 | 0 |
| Human fears | Hmnfrs | 3 | 1 | 1 |
| Human health issues | Hmnhli | 5 | 3 | 40 |
| Negative impact on beneficial organisms | Ngiobo | 11 | 0 | 0 |
| Spreading diseases | Sprdnd | 1 | 1 | 4 |

Table 4: ESP nodes: degrees and realised evidence (ES/EDS links and paper weights in brackets).

| Node | Abbr. | Pred. degree (ES; EDS) | Real. degree (ES; EDS) | Real. papers (ES; EDS) |
| --- | --- | --- | --- | --- |
| Amphibians | Amphbn | 3 (ES 1; EDS 2) | 1 (ES 1; EDS 0) | 2 (ES 2; EDS 0) |
| Bacteria | Bacter | 4 (ES 3; EDS 1) | 4 (ES 3; EDS 1) | 106 (ES 83; EDS 23) |
| Beetles | Beetls | 9 (ES 6; EDS 3) | 7 (ES 6; EDS 1) | 102 (ES 88; EDS 14) |
| Birds | Birds | 11 (ES 7; EDS 4) | 5 (ES 4; EDS 1) | 95 (ES 78; EDS 17) |
| Bugs and aphids etc. | Baaet. | 5 (ES 2; EDS 3) | 4 (ES 2; EDS 2) | 75 (ES 60; EDS 15) |
| Butterflies and moths | Bttram | 4 (ES 3; EDS 1) | 4 (ES 3; EDS 1) | 68 (ES 46; EDS 22) |
| Earthworms | Erthwr | 3 (ES 2; EDS 1) | 0 (ES 0; EDS 0) | 0 (ES 0; EDS 0) |
| Flies and mosquitos | Flsanm | 3 (ES 1; EDS 2) | 3 (ES 1; EDS 2) | 46 (ES 40; EDS 6) |
| Fungi | Fungi | 7 (ES 4; EDS 3) | 6 (ES 4; EDS 2) | 310 (ES 285; EDS 25) |
| Mammals | Mammls | 14 (ES 7; EDS 7) | 6 (ES 4; EDS 2) | 28 (ES 20; EDS 8) |
| Microsporidia | Mcrspr | 2 (ES 1; EDS 1) | 1 (ES 1; EDS 0) | 1 (ES 1; EDS 0) |
| Mites | Mites | 3 (ES 1; EDS 2) | 2 (ES 1; EDS 1) | 41 (ES 39; EDS 2) |
| Molluscs | Mllscs | 2 (ES 1; EDS 1) | 1 (ES 0; EDS 1) | 2 (ES 0; EDS 2) |
| Nematods | Nemtds | 3 (ES 2; EDS 1) | 3 (ES 2; EDS 1) | 53 (ES 40; EDS 13) |
| Reptiles | Reptls | 3 (ES 1; EDS 2) | 1 (ES 1; EDS 0) | 1 (ES 1; EDS 0) |
| Spiders | Spidrs | 2 (ES 1; EDS 1) | 1 (ES 1; EDS 0) | 99 (ES 99; EDS 0) |
| Springtails | Sprngt | 4 (ES 3; EDS 1) | 4 (ES 3; EDS 1) | 8 (ES 7; EDS 1) |
| Viruses | Viruss | 1 (ES 0; EDS 1) | 1 (ES 0; EDS 1) | 3 (ES 0; EDS 3) |
| Wasps, bees and ants | Ws,baa | 4 (ES 2; EDS 2) | 2 (ES 2; EDS 0) | 104 (ES 104; EDS 0) |
| Wild plants | Wldpln | 16 (ES 13; EDS 3) | 2 (ES 2; EDS 0) | 10 (ES 10; EDS 0) |

Figure 11: Distribution of degree loss (predicted minus realised) for ES vs EDS nodes. EDS nodes (red) show a rightward shift, indicating greater loss of connectivity relative to ES nodes (green).

### Network projections

Table 5: ES/EDS bundle structure in predicted and realised networks. For each ecosystem service (ES) or disservice (EDS), the number of ES and EDS neighbours in the projection network (bundled through shared ESPs) and the total number of ESPs mediating those bundles are shown separately for predicted and realised networks.

|  | | Predicted | | | | Realised | | | |
| --- | --- | --- | --- | --- | --- | --- | --- | --- | --- |
| ES/EDS | Type | Links (ES) | Links (EDS) | ESPs (ES) | ESPs (EDS) | Links (ES) | Links (EDS) | ESPs (ES) | ESPs (EDS) |
| Damage on crop | EDS | 18 | 8 | 52 | 19 | 9 | 3 | 30 | 4 |
| Ecosystem engineering | EDS | 8 | 6 | 9 | 7 | 0 | 0 | 0 | 0 |
| Faciliation of the spread of the weeds | EDS | 10 | 6 | 20 | 11 | 0 | 0 | 0 | 0 |
| Facilitation of pest dispersal | EDS | 7 | 6 | 14 | 9 | 0 | 0 | 0 | 0 |
| Habitat and food for pests | EDS | 13 | 2 | 13 | 2 | 0 | 0 | 0 | 0 |
| Human fears | EDS | 7 | 6 | 9 | 8 | 4 | 1 | 4 | 1 |
| Human health issues | EDS | 18 | 7 | 28 | 11 | 6 | 1 | 8 | 2 |
| Negative impact on beneficial organisms | EDS | 11 | 7 | 33 | 19 | 0 | 0 | 0 | 0 |
| Spreading diseases | EDS | 2 | 2 | 2 | 2 | 2 | 1 | 2 | 1 |
| Aesthetics | ES | 15 | 8 | 26 | 15 | 2 | 1 | 2 | 1 |
| Biopesticide | ES | 12 | 3 | 12 | 3 | 0 | 0 | 0 | 0 |
| Climate and weather regulation | ES | 12 | 3 | 12 | 3 | 0 | 0 | 0 | 0 |
| Cultural practices | ES | 14 | 8 | 24 | 14 | 0 | 0 | 0 | 0 |
| Decomposition | ES | 8 | 4 | 14 | 10 | 8 | 2 | 13 | 6 |
| Detoxification | ES | 14 | 3 | 14 | 4 | 2 | 1 | 2 | 1 |
| Fibre | ES | 12 | 3 | 12 | 3 | 0 | 0 | 0 | 0 |
| Food provision | ES | 17 | 8 | 29 | 19 | 7 | 3 | 12 | 6 |
| Fuel | ES | 12 | 3 | 12 | 3 | 0 | 0 | 0 | 0 |
| Pest control | ES | 17 | 9 | 37 | 35 | 7 | 4 | 20 | 11 |
| Pharmaceutical | ES | 12 | 3 | 12 | 3 | 0 | 0 | 0 | 0 |
| Pigment | ES | 16 | 5 | 17 | 6 | 5 | 1 | 5 | 1 |
| Pollination | ES | 7 | 5 | 8 | 8 | 7 | 2 | 8 | 4 |
| Recreation | ES | 14 | 8 | 24 | 14 | 0 | 0 | 0 | 0 |
| Seed dispersal | ES | 9 | 7 | 17 | 14 | 6 | 2 | 11 | 4 |
| Soil functions | ES | 4 | 3 | 8 | 6 | 4 | 2 | 7 | 4 |
| Timber | ES | 12 | 3 | 12 | 3 | 0 | 0 | 0 | 0 |
| Weed control | ES | 9 | 8 | 18 | 17 | 6 | 3 | 12 | 6 |

Table 6: ESP projection structure in predicted and realised networks. For each ecosystem service provider (ESP), the degree in the ESP projection network (number of other ESPs linked through shared services/disservices) and the number of ES and EDS that are shared with at least one other ESP are shown separately for predicted and realised networks.

|  | Predicted | | | Realised | | |
| --- | --- | --- | --- | --- | --- | --- |
| ESP | Linked ESPs | Shared ES | Shared EDS | Linked ESPs | Shared ES | Shared EDS |
| Amphibians | 13 | 2 | 1 | 13 | 1 | 0 |
| Bacteria | 8 | 1 | 3 | 5 | 2 | 1 |
| Beetles | 19 | 3 | 6 | 18 | 5 | 1 |
| Birds | 18 | 4 | 7 | 17 | 4 | 1 |
| Bugs and aphids etc. | 18 | 2 | 2 | 17 | 2 | 1 |
| Butterflies and moths | 14 | 1 | 3 | 13 | 2 | 1 |
| Earthworms | 14 | 1 | 2 | 0 | 0 | 0 |
| Flies and mosquitos | 15 | 2 | 1 | 13 | 1 | 2 |
| Fungi | 19 | 3 | 4 | 18 | 4 | 2 |
| Mammals | 19 | 7 | 7 | 17 | 4 | 1 |
| Microsporidia | 13 | 1 | 1 | 13 | 1 | 0 |
| Mites | 18 | 2 | 1 | 17 | 1 | 1 |
| Molluscs | 13 | 1 | 1 | 11 | 0 | 1 |
| Nematods | 19 | 1 | 2 | 18 | 2 | 1 |
| Reptiles | 13 | 2 | 1 | 13 | 1 | 0 |
| Spiders | 13 | 1 | 1 | 13 | 1 | 0 |
| Springtails | 19 | 1 | 3 | 18 | 3 | 1 |
| Viruses | 13 | 1 | 0 | 11 | 0 | 1 |
| Wasps, bees and ants | 15 | 2 | 2 | 15 | 2 | 0 |
| Wild plants | 19 | 2 | 7 | 14 | 2 | 0 |

Figure 12: Predicted network - Projection 1

Figure 13: Predicted network - Projection 2

Figure 14: Filtered network with weights - Projection 1

Figure 15: Filtered network with weights - Projection 2

Figure 16: Filtered network, no weights - Projection 1

Figure 17: Filtered network, no weights - Projection 2
